# Fast and Accurate *in silico* Antigen Typing with Kaptive 3

**DOI:** 10.1101/2025.02.05.636613

**Authors:** Thomas D. Stanton, Marit A.K. Hetland, Iren H. Löhr, Kathryn E. Holt, Kelly L. Wyres

**Author notes:** **Correspondence**: Dr. Tom Stanton, Dr. Kelly Wyres, The Alfred and Monash University, Level 2, The Burnet Institute, 85 Commercial Road, Melbourne VIC 3004, Australia. E.

## Abstract

Surface polysaccharides are common antigens in priority pathogens and therefore attractive targets for novel control strategies such as vaccines, monoclonal antibody and phage therapies. Distinct serotypes correspond to diverse polysaccharide structures that are encoded by distinct biosynthesis gene clusters, e.g. the *Klebsiella pneumoniae* species complex (KpSC) K- and O-loci encode the synthesis machinery for the capsule (K) and outer-lipopolysaccharides (O), respectively. We previously presented Kaptive and Kaptive 2, programs to identify K and O-loci directly from KpSC genome assemblies (later adapted for *Acinetobacter baumannii*), enabling sero-epidemiological analyses to guide vaccine and phage therapy development. However, for some KpSC genome collections, Kaptive (v≤2) was unable to type a high proportion of K-loci. Here we identify the cause of this issue as assembly fragmentation, and present a new version of Kaptive (v3) to circumvent this problem, reduce processing times and simplify output interpretation.

We compared the performance of Kaptive v2 and Kaptive v3 for typing genome assemblies generated from subsampled Illumina read sets (decrements of 10x depth), for which a corresponding high quality completed genome was also available to determine the ‘true’ loci (n=549 KpSC, n=198 *A. baumannii*). Both versions of Kaptive showed high rates of agreement to the matched true locus among ‘typeable’ locus calls (≥96% for ≥20x read depth), but Kaptive v3 was more sensitive, particularly for low depth assemblies (at <40x depth, v3 ranged 0.85-1 vs v2 0.09-0.94) and/or typing KpSC K-loci (e.g. 0.97 vs 0.82 for non-subsampled assemblies). Overall, Kaptive v3 was also associated with a higher rate of optimal outcomes i.e. loci matching those in the reference database were correctly typed and genuine novel loci were reported as untypeable (73-98% for v3 vs 7-77% for v2 for KpSC K-loci).

Kaptive v3 was >1 order of magnitude faster than Kaptive v2 making it easy to analyse thousands of assemblies on a desktop computer, facilitating broadly accessible in silico serotyping that is both accurate and sensitive. The Kaptive v3 source code is freely available on GitHub (https://github.com/klebgenomics/Kaptive), and has been implemented in Kaptive Web (https://kaptive-web.erc.monash.edu).

## Introduction

Bacterial surface polysaccharides such as capsules and lipopolysaccharides (LPS) play a key role in protection from host immune systems, phagocytosis, bacteriophage predation and desiccation [1,2]. Many act as primary phage receptors and are highly immunogenic, making them major targets for vaccines, phage and monoclonal antibody therapies. However, therapeutic design is complicated by broad structural diversities that result in a differential phage susceptibility and a wide variety of immunologically distinct serotypes [3–5].

Glycoconjugate vaccines that target a subset of known serogroups/serotypes have played a pivotal role in preventing pneumococcal and meningococcal infections, providing an exemplar for effective use of multi-valent formulations [6]. With the growing burden of antimicrobial resistance, there is increasing interest in the development of novel control strategies targeting other species, particularly priority pathogens such as *Klebsiella pneumoniae* and *Acinetobacter baumannii* [7–9]. However, there is a lack of broadly accessible serological typing schemes to support therapeutic design.

In the *K. pneumoniae* Species Complex (KpSC), capsule (K) and outer-LPS (O) antigens are produced through biosynthetic pathways encoded by corresponding K- and O-locus gene clusters [10,11]. Similarly, in *A. baumannii* the capsule (K) and outer-core (OC) antigens are encoded by the K- and OC-loci, respectively. Each locus comprises a set of conserved export and assembly machinery genes, along with a unique set of glycosidic linkage and modification genes that result in unique polysaccharide structures. Therefore, K- and O/OC-types can be predicted directly from the gene content of the cluster [12–14]. With this knowledge and the continued growth of whole-genome sequencing, we have new opportunities to investigate capsule and LPS diversity and epidemiology, and prioritise polysaccharide variants for novel control strategies [15].

We previously reported 134 unique K-loci from KpSC genomes and developed Kaptive, a tool to rapidly type K-loci from bacterial genome assemblies [16]. Since Kaptive’s release, 52 additional KpSC K-loci have been reported [17], as well as an O-locus database [18], plus K- and OC-locus databases for *A. baumannii* [19] (distributed via GitHub alongside the Kaptive code at https://github.com/klebgenomics/Kaptive). Kaptive-compatible *Vibrio parahaemolyticus* K- and O-locus databases are hosted in a third-party repository [20], and a mixture of partial and complete databases have been described for other organisms [21–25]. In Kaptive v0.4.0 we implemented logic to account for modification of the KpSC O2 antigen based on specific genes outside of the O-locus [17]. This logic was generalised in Kaptive v2, when we added an explicit phenotype prediction column in the Kaptive output. It was later extended to the *A. baumannii* K-locus database after the finding that the presence of a phage-encoded Wzy protein resulted in altered polysaccharide structures [17,26].

The Kaptive v≤2 algorithm has two main steps. First BLASTn is used to align the full-length reference locus nucleotide sequences against the input assembly contigs, and the best-match reference is chosen as the one with highest overall alignment coverage. Then tBLASTn is used to align translated protein sequences from each reference against the input assembly, mark genes inside and outside of the assembly locus region and report any expected genes that are missing and/or any unexpected genes that are present (i.e. genes that are not present in the best-match reference). tBLASTn is additionally used to search for genes outside the locus that are known to impact phenotype.

Kaptive v≤2 report 6-tier confidence scores (**Table S1**) that were developed based on the logic of locus definitions and our working experiences with KpSC draft genome assemblies; however, no systematic testing was completed. The scores penalise missing and extra genes within the locus region of the assembly, and fragmented loci (on the basis that we cannot be sure that we have detected the full complement of genes that may be present on other assembly contigs or erroneously missing from the assembly). These scores were intended to guide users with interpretation and follow-up investigations appropriate to their specific use-case; however, in practice we have observed that most users either simply follow our baseline recommendation to exclude ‘Low’ and ‘None’ confidence scores or ignore the confidence scores entirely. Additionally, some datasets have a high rate of ‘Low’ and ‘None’ confidence scores, rendering large amounts of data unusable for sero-epidemiological analysis, e.g. 36.8% (n=121/329) in a study of invasive KpSC isolates from South and South East Asia, and 32.6% (n=84/258) in a study of *K. pneumoniae* neonatal sepsis isolates from seven distinct countries [27,28].

Here we show that low Kaptive v≤2 confidence scores are driven by assembly fragmentation, which often results from a failure to incorporate low-GC regions of the K-locus into Illumina sequencing libraries. We present an updated version of Kaptive (v3), with several performance enhancements and a simplified confidence scoring system to address the limitations associated with fragmented assemblies. We also perform a systematic comparison and show that Kaptive v3 is highly accurate, more sensitive and faster than Kaptive v2.

## Materials and Methods

### Kaptive v3 Specifications

Kaptive v3 is a Python application (v3.9) that builds upon the existing open-source code developed for Kaptive v≤2 [16,17] (available at https://github.com/klebgenomics/Kaptive). While the prior versions used a combination of BLASTn and tBLASTn [29] for sequence search, Kaptive v3 uses minimap2 [30] to search for gene nucleotide sequences, of which a non-overlapping subset are translated and pairwise-aligned to the respective references with Biopython (v1.83) [31]. We chose to remove the BLAST+ dependency because it is comparatively slow, and some versions are subject to random crashes during multi-threaded tBLASTn. DNA features viewer [32] is used to optionally generate locus images.

We additionally refactored the Kaptive source code from a single command line interface (CLI) script into a more efficient, user-friendly Python package with an API, allowing Kaptive to be imported as a module and used in other programs. The main mode of operation, *in silico* serotyping of genome assemblies, is executed by the “typing pipeline” and implemented via the “assembly” CLI mode. Kaptive databases are parsed from GenBank files into Database objects, which hold the sequence information in memory. These objects have formatting methods allowing them to be converted into other biological text formats (e.g. nucleotide or protein sequence fasta files), implemented via the “extract” CLI mode. The new “convert” CLI mode allows the conversion of the typing results in JSON format (now using the more efficient JSON lines format) into other biological text formats to facilitate downstream investigations.

The new locus typing approach is described in detail in Results. The Kaptive v3 source code is published under GNU General Public License v3.0 and available at https://github.com/klebgenomics/Kaptive. It has been implemented in the web-based graphical user interface tools Kaptive Web v1.3.0 (https://kaptive-web.erc.monash.edu/) and Pathogenwatch (https://pathogen.watch/) for K- and O/OC-locus typing of KpSC and *A. baumannii.* It has also been implemented in the CLI tool Kleborate v3 (https://github.com/klebgenomics/kleborate), and the Bactopia CLI pipeline [33], for KpSC K- and O-locus typing.

### Test Dataset

To test the accuracy of Kaptive v3 locus typing from draft genome sequences, we sourced collections of high-quality completed genome assemblies (i.e. circularised via hybrid assembly) with corresponding short-reads, to generate a dataset where we could compare locus calls from draft short-read-based assemblies to ‘ground truth’ locus calls derived from the matched completed genome. We utilised 549 diverse KpSC genomes representing clinical and gut carriage isolates collected from humans and animals in Norway [34]. For *A. baumannii*, we compiled a collection of 198 completed genomes deposited in the National Center for Biotechnology Information (NCBI) Assembly database, which had corresponding paired-end Illumina reads deposited in the SRA database (**Table S2**). All high-quality completed assemblies were annotated with Bakta v1.9.2 [35], and preliminary K-, O- and OC-loci were assigned using the best BLASTn coverage approach implemented in Kaptive v2.0.9 [17,19,36]. The loci were extracted from the corresponding Bakta GenBank annotations and visually inspected with Clinker v0.0.29 [37] to confirm ground truth calls. As per the locus definition rules, loci were confirmed as the best match if they comprised the complete set of genes present in the reference locus and no additional genes (excluding transposases and ignoring pseudogenes). Distinct genes were defined based on the species-specific translated sequence identity thresholds; 82.5% for KpSC, 85% for *A. baumannii* [16,19]. Loci with ≥1 polysaccharide-specific gene missing and/or ≥1 additional gene (excluding transposases) compared to the best matching reference were marked as novel. Loci with ≥1 insertion sequence (IS) insertion, but which otherwise contained the same gene set as the best match reference, were marked as IS variants. Loci that were missing ≥1 core assembly machinery gene but otherwise contained the same gene set as the best match reference were marked as deletion variants (presumed to represent isolates that are unable to produce and/or export the relevant polysaccharide).

The finalised KpSC ground truth collections captured 96 distinct K-loci (plus 9 genomes with novel loci, 7 with deletion and 79 with IS variants) and 11 distinct O-loci (plus 10 genomes with novel, 1 deletion and 16 IS variants). The *A. baumannii* ground truth collection captured 45 distinct K-loci (plus 2 genomes with novel and 14 IS variants) and 13 distinct OC-loci (plus 1 genome with a novel locus and 41 IS variants).

We next generated increasingly fragmented or “low-quality” draft assemblies for each genome by randomly subsampling the corresponding Illumina short-reads from 100x-10x mean read depth in decrements of 10 with Rasusa v0.7.1 [38] (using index files generated with Samtools v1.9 [39]), and assembling them with Unicycler v0.5.0, with the “--depth_filter” flag set to 0. For the 549 KpSC genomes, we obtained the following draft assemblies: n=161 100x depth, n=176 90x, n=167 80x, n=200 70x, n=230 60x, n=223 50x, n=442 40x, n=549 (all genomes) at 30x, 20x, 10x, and without subsampling. Note these sample sizes were constrained by the estimated read depths of the non-subsampled data. For the 198 *A. baumannii* genomes, we obtained n=198 draft assemblies for all subsampling depths and without subsampling.

### Illumina Read Coverage and GC content

We used Biopython (v1.83) to determine the GC content of K and O/OC-locus gene ORFs for each of the complete genomes from the Bakta GenBank annotations, along with the GC content of their respective chromosome. Here, we define GC as (ΣG + ΣC) / sequence length, and GC difference as |gene GC – chromosome GC| where chromosome GC is ∼0.57 for KpSC [40] and ∼0.39 for *A. baumannii*. To label each gene as core or variable, we clustered the translated amino acid sequences for each locus database using the MMSeqs2 (v15-6f452) “easy-cluster” command [41]. We defined core and variable genes as those belonging to clusters represented in ≥75% and <75% reference loci respectively. We mapped the Illumina short-reads to the corresponding completed genome assemblies with minimap2 (v2.2.0) [30] using the parameters “-c -x sr”. The resulting PAF format alignments were converted to BED format with the paftools “splice2bed” command (see minimap2) and read coverage was calculated with the bedtools (v2.31.0) “coverage” command [42] using the Bakta GFF annotation files.

### Kaptive Performance Comparisons and Benchmarking

We compared the typing performance of Kaptive v2.0.9 and Kaptive v3.0.0.b5 for all assembly types. We defined agreement as the percentage of correct and typeable (true positive) calls, and sensitivity as true positive / (true positive + false negative), where false negatives were defined as correct but untypeable calls. Typeable loci were defined as those with confidence score ‘Typeable’ (Kaptive v3) and any of ‘Good’, ‘High’, ‘Very high’ or ‘Perfect’ (Kaptive v2). We report the rates of typeability (regardless of agreement), agreement and sensitivity across both Kaptive versions. We do not report specificity or accuracy as per the standard statistical definitions due to the difficulty in defining true negative outcomes: While an incorrect call marked as ‘untypeable’ could be considered a true negative outcome, the vast majority do not represent a bona fide true negative because there are very few genuine novel (untypeable) loci in the test datasets. As a consequence, we observed a high number of incorrect and ‘untypeable’ calls for assemblies that harbour loci represented in the reference database that should therefore be typeable. These calls resulted in inflated true negative counts, and misleading specificity/accuracy estimates. Therefore, we instead report the rate of optimum outcomes as the sum of the percentages: i) of assemblies harbouring a locus with a true match in the reference database, that were reported correctly and marked as ‘typeable’; and ii) of assemblies harbouring a genuine novel locus, reported as ‘untypeable.’

Finally, we benchmarked runtime performance between Kaptive v2.0.9 and Kaptive v3.0.0.b5 on a desktop computer with a single ARM Apple M2 8 Core CPU. Runtime was measured using the “time” utility (Zsh built-in) across both the K and O/OC-locus databases for each completed KpSC and *A. baumannii* assembly (n=1494) using the commands: “kaptive assembly <DB> <ASSEMBLY> -o /dev/null” for Kaptive 3 and “python kaptive.py -k <DB> -a <ASSEMBLY> --threads 8 -o /dev/null” for Kaptive 2. Both versions used 8 threads for alignment (noting that Kaptive 3 will automatically default to the maximum number of available CPUs or cap out at 32), and the Kaptive v2 assembly BLAST+ databases were cleared after each run so that database construction time was included in each benchmark.

Python v3.10 and R v4.4.1 were used for scripts and statistical analyses unless otherwise stated.

## Results

### Illumina Sequence Coverage Influences Locus Fragmentation and Kaptive v2 Typeability

We explored the Kaptive v2 calls of short-read draft assemblies (no read subsampling), excluding those harbouring novel/deletion variants as determined by inspection of the complete genome sequence (considered the ground truth). For KpSC K-loci we found that 25% (n=134/533) were assigned confidence ‘Low’ or ‘None’ and would therefore be considered ‘untypeable’, whereas only 2% of O-loci (n=12/538) had these confidence values. For *A. baumannii* 8% (n=15/196) and 4% (n=7/197) K- and OC-loci were assigned these values, respectively. We also noted that many of the KpSC K-loci (48%, n=257/533) were fragmented over contigs and a similar number (48%, n=254/533) were lacking one or more expected genes, with both events usually co-occurring in the same assemblies. However, among the KpSC O-loci and *A. baumannii* K/OC-loci, ≤25% were fragmented and/or missing genes, respectively.

Most of the missing K-locus genes in the Illumina-only (non-subsampled) KpSC assemblies were involved in sugar processing (n=923/1088, 85%). These genes had a mean absolute GC difference of 0.19 (where GC difference is defined as the GC of a gene minus the GC value for the chromosome). This mean absolute difference was over double the mean absolute GC difference of missing genes in the O-locus (0.08) (**Figure S1a, Table S3**). We speculated that these genes may be missing or fragmented in the assemblies due to GC-dependent sequencing dropout i.e. when a region of a genome is not captured in the Illumina read data because its GC content differs substantially from the genome mean value for which the library preparation protocol is optimised. We therefore tested for an association between GC difference and Illumina sequencing coverage of all K-locus genes stratified by their prevalence among K-loci (core vs variable) and the library preparation kit (Nextera Flex vs Nextera XT). There was a significant association for Illumina reads prepared with the Nextera XT kit (p<0.001, W=825411.5; using a Wilcoxon rank sum test with continuity correction), which was not apparent for those prepared with the newer Nextera Flex kit (p=0.212) (**Figure 1**). The impact was greatest for variable genes i.e. the capsule specific sugar processing genes (**Figure 1**), which are known to be associated with comparatively greater GC divergence from the chromosome [16].

**Figure 1:**
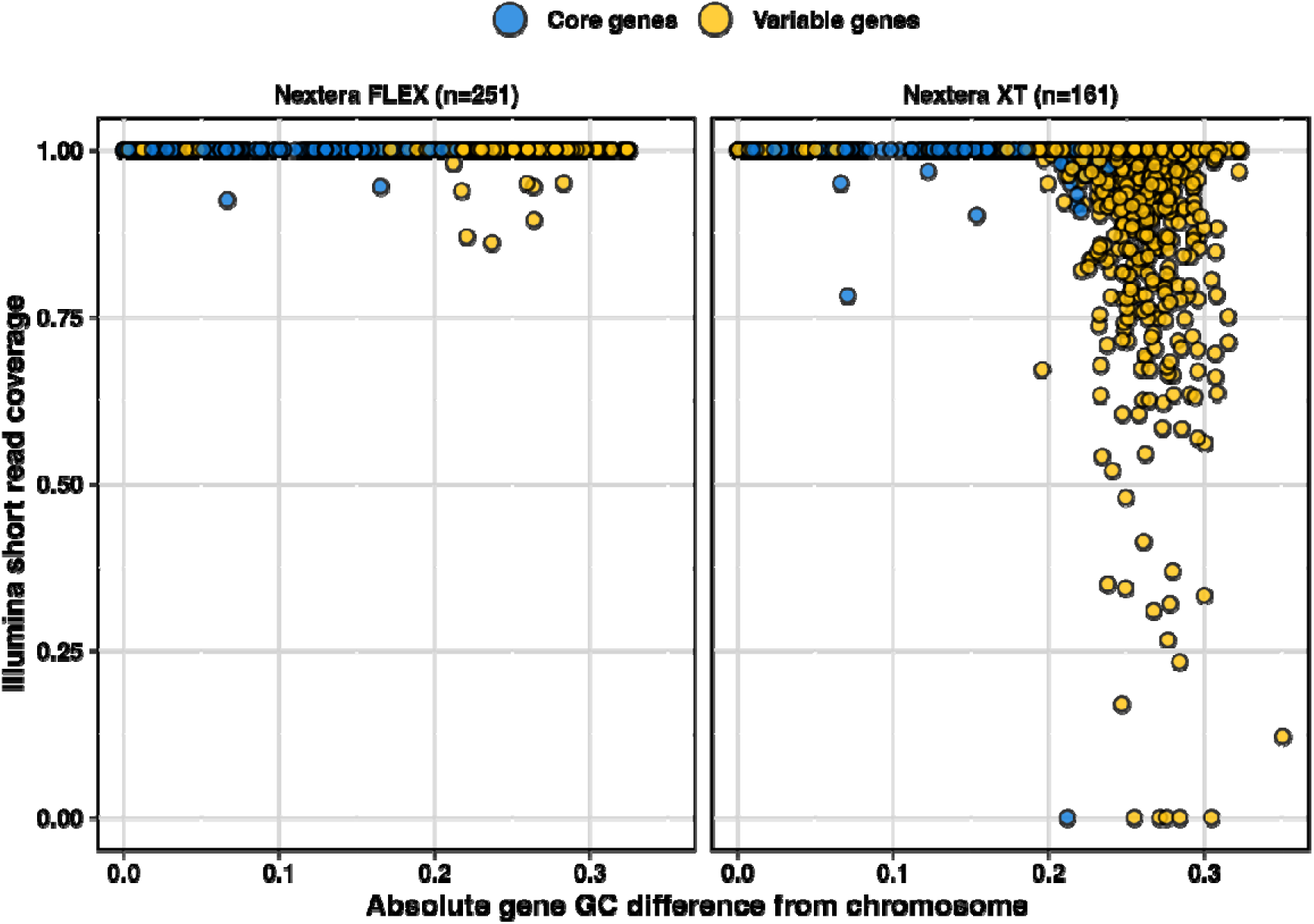
Illumina sequencing coverage vs absolute GC difference from the chromosome. K-locus genes from matched completed KpSC assemblies are stratified by the library preparation chemistry and coloured by prevalence among reference loci, as indicated. Core genes represent those encoding the conserved synthesis and export machinery; variable genes are those involved in sugar processing and antigenic variation.

Similar GC-content-dependent coverage bias has been reported previously for sequencing libraries prepared with the Nextera XT kit due to variation in tagmentation site cleavage efficiency [43]. This bias likely results in non-amplification of K-locus gDNA during Illumina library preparation, and subsequent data loss during sequencing. This data loss perturbs De Bruijn-graph-based assembly in this region of the chromosome, resulting in K-loci that are “broken” over contigs (i.e. fragmented into multiple pieces), fuelling the low K-locus typeability for KpSC. Notably, absolute GC differences were much lower for KpSC O and *A. baumannii* K/OC-locus genes (**Figure S1a**) and there was comparatively minimal sequencing drop-out (**Figure S1b**).

### Locus typing with Kaptive v3

To overcome the typeability issues resulting from fragmented assemblies, Kaptive 3 uses a two-stage gene-first locus scoring algorithm to maximise information entropy from nucleotide alignments (**Figure 2**).

**Figure 2:**
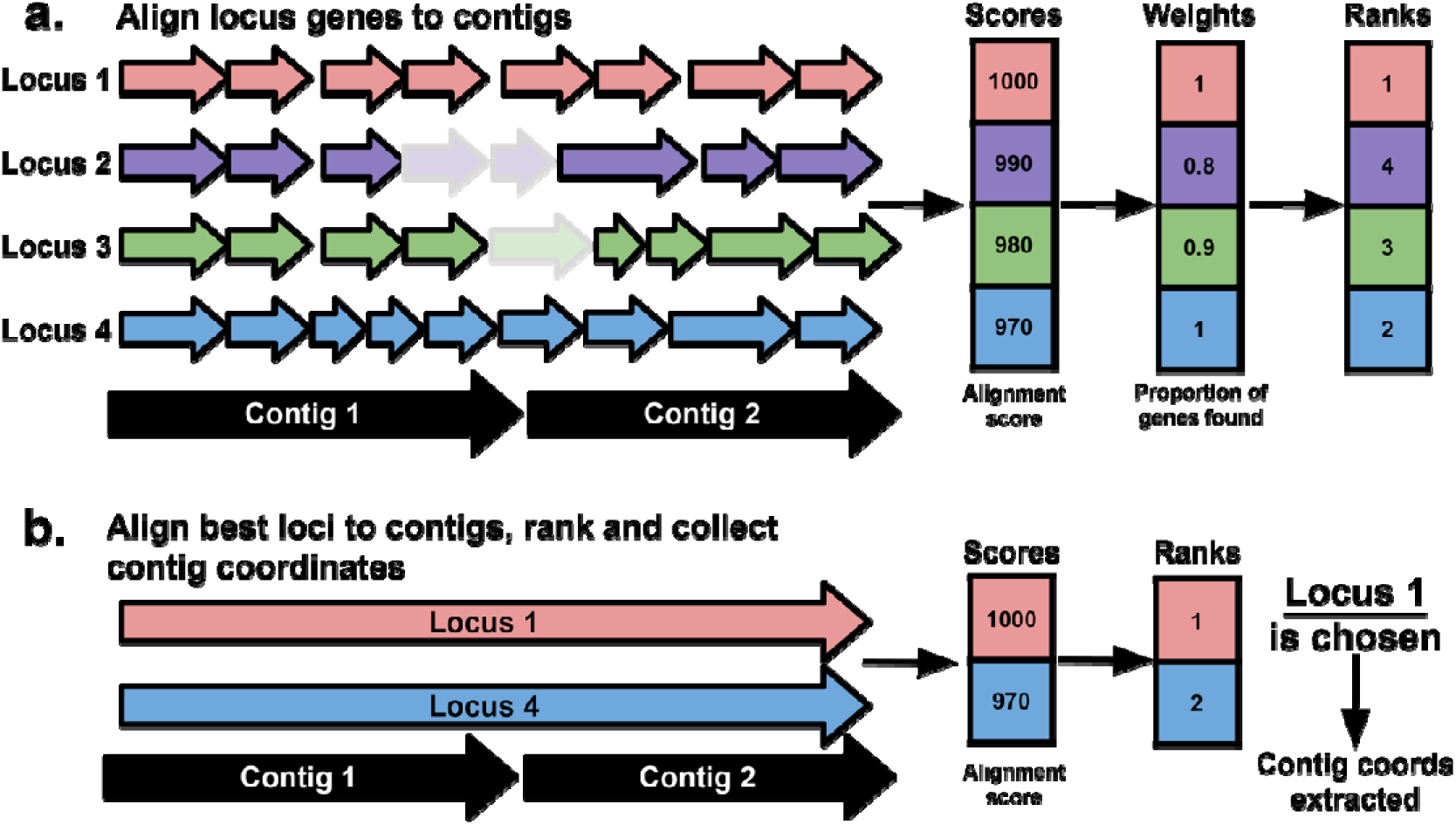
Overview of the Kaptive 3 locus scoring algorithm stages. (a) Reference locus genes are aligned to input assembly contigs with minimap2, alignment metrics are summed, weighted and ranked. The default alignment and scoring metrics are shown; alignment score and proportion of locus genes found, respectively. Shading indicates genes with (dark) vs without (light) alignments. (b) The full nucleotide sequences of the top-ranking loci from the first stage are aligned to the contigs to achieve further resolution and determine the locus coordinates in the input assembly.

In the first stage, minimap2 is used to align all genes from all reference loci to the input assembly contig sequences. A score is calculated for each locus by summing the specified alignment metric for all of the corresponding genes (either alignment score, number of matching bases, number of aligned bases or query length). The alignment scores are weighted by the specified weighting metric (number of genes found, number genes expected, proportion of genes found or total length of each reference locus) and each locus is ranked (**Figure 2a**). The default alignment and weighting metrics are “alignment score” and “proportion of genes found” selected as those that resulted in the highest proportion of locus calls matching the ground truth for the test dataset (**Table S4**).

In the second stage, the full-length nucleotide sequences of the top-ranking reference loci are fully aligned to the assembly, and the best-match locus is identified based on the best alignment score (**Figure 2b**). This second step provides coordinates of the locus within the assembly and allows Kaptive to distinguish between closely related loci with highly similar gene content such as OCL1 and OCL9 in *A. baumannii,* that differ by the presence of a single gene [36]. The number of top-ranking loci fully aligned to the assembly can be specified by the user using the “--n-best” parameter, with a default of 2 (based on observations of pairs of highly similar *A. baumannii* loci; however, for databases without pairs of similar loci this can be set to 1 to simply retrieve the coordinates of the best locus from the first stage; or for databases with groups of highly similar loci the parameter can be increased).

Once the best matching reference locus has been identified, the original gene alignments are culled to remove overlapping alignments corresponding to orthologous genes from different reference loci, with a preference to retain those associated with the best matching locus. The nucleotide sequences of the remaining gene alignments are extracted from the input assembly, and if the gene coordinates overlap the full-length reference locus alignment coordinates, the gene is annotated as part of the locus. Each gene nucleotide sequence is translated and aligned to the respective reference locus protein using the Smith-Waterman algorithm to determine variation in amino-acid space [44]. Protein identity and coverage compared to the references are reported in the Kaptive output.

Extra-locus genes, i.e. genes outside of the K, O and OC-loci that are known to impact polysaccharide phenotypes, are detected and reported as described above. Additionally, we have implemented a new phenotypic prediction logic which updates the predicted polysaccharide type based on the final intact gene content of the locus, using known phenotype-genotype patterns from the literature, e.g. truncation of the WcaJ or WbaP initiating glycosyltransferase proteins encoded by KpSC K-loci results in a capsule-null (acapsular) phenotype [45]. Files containing the specific logic for each species and locus can be found in the Kaptive reference database directory in the Kaptive git repository, and are further described in the Kaptive Database documentation (https://kaptive.readthedocs.io/en/latest/Databases.html).

### Kaptive v3 confidence scores

We redefined Kaptive’s confidence criteria with consideration of the locus definition rules (i.e. that each locus represents a unique set of genes defined at a given minimum translated identity threshold) and to optimise the balance of correct vs incorrect (un)typeable calls, especially for highly fragmented assemblies (**Table 1, Figure S2**). We also sought to make the confidence calls easier to interpret and have simplified the confidence tiers to explicitly state “Typeable” or “Untypeable”.

**Table 1:** Kaptive 3 confidence score definitions. When a locus is identified in a single contiguous piece within the input assembly, we apply strict criteria to define it as a ‘Typeable’ match to the reference locus, i.e. it contains the same set of genes (note that pseudogenes are counted here). When a locus is not contiguous in the input assembly, we allow greater flexibility to account for sequencing drop-out and misalignment of partial gene sequences.

| Confidence | Fragmented | # Genes below identity threshold | Expected genes found (%) | # Extra genes |
| --- | --- | --- | --- | --- |
| Typeable | No | 0 | 100 | 0 |
|  | Or if locus is fragmented |  |  |  |
|  | Yes | 0 | ≥50 | ≤1 |
| Untypeable | Does not meet the above criteria |  |  |  |

### Kaptive v3 is Highly Sensitive and Accurate

Kaptive v3 reported a typeable result for a greater number of assemblies than Kaptive v2 for all databases and assembly types (**Figure S3, Table S5**). Among the assemblies that were classified as ‘typeable’, percent agreement was high for both Kaptive v2 and v3 (**Figure 3, Table S5**) i.e. the vast majority of results were reported as the correct locus (≥91% for all databases and read depths, ≥96% for read depths >20x). Misidentified loci primarily comprised IS variants and a minority of genuinely novel loci that were not represented in the reference databases (see **Supplementary Results**). Among assemblies that were classified as ‘untypeable,’ rates of agreement were varied i.e. many of the reported best-match loci did not match the ground truth, particularly for Kaptive v2 (see **Supplementary Results, Figure S4, Table S5**). In such instances the ‘untypeable’ classification is appropriate, although the overall outcome is sub-optimal (see below).

**Figure 3:**
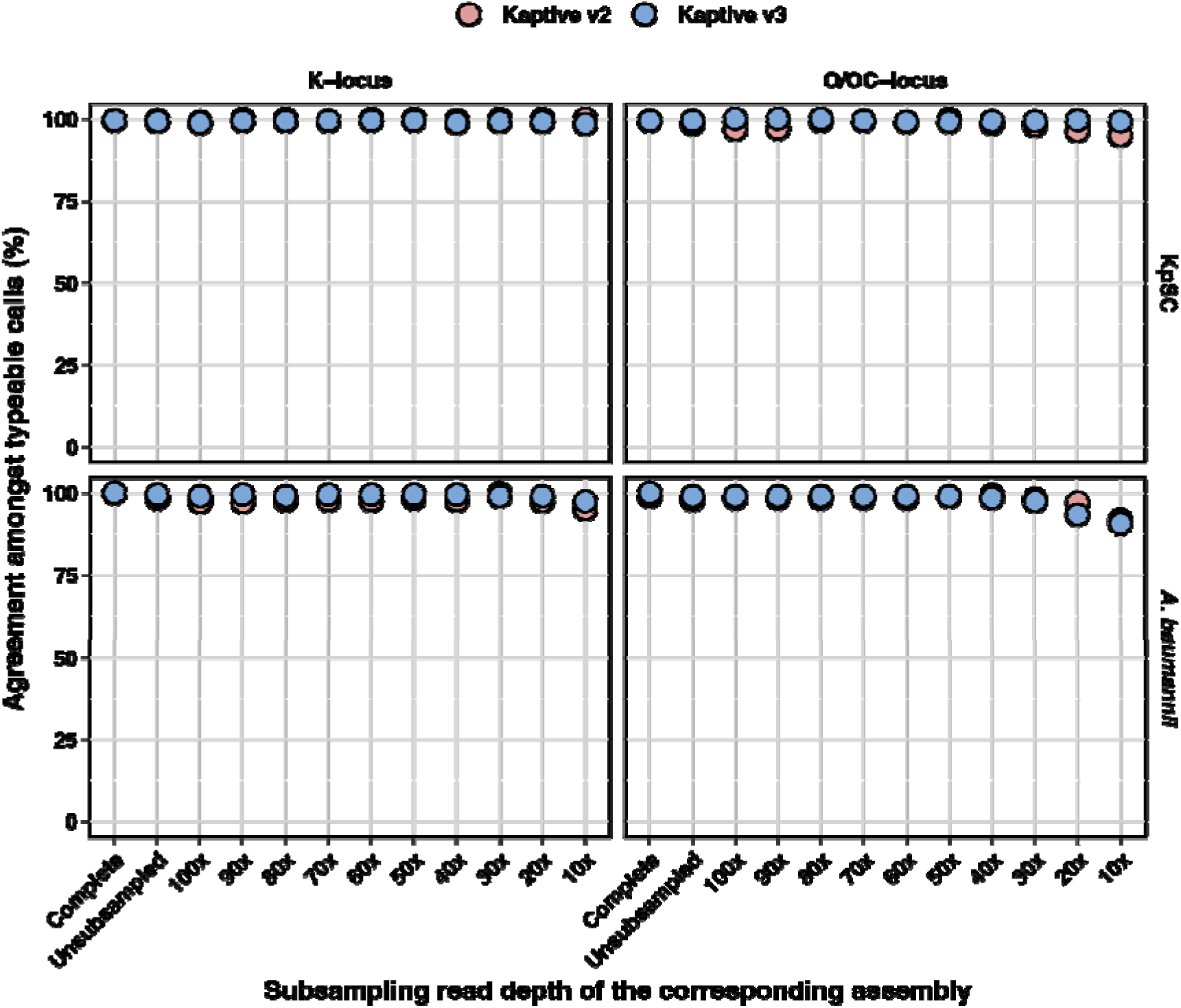
Percent agreement amongst ‘typeable’ Kaptive locus calls. Data are stratified by species (rows), database (columns) and assembly subsampling group, and coloured by Kaptive version as indicated. Assembly subsampling groups are arranged on the X-axis in descending order of subsampling read depth, starting with the complete (hybrid) and non-subsampled Illumina-only assemblies, then subsampling the Illumina-only reads at the stated increments.

Sensitivity reflects the proportion of assemblies carrying true matches to loci in the reference database that are correctly identified *and* reported as ‘typeable.’ Consistent with the differences in typeability, sensitivity was notably higher for Kaptive v3 than for Kaptive v2, particularly for low depth assemblies (<40x depth, sensitivity range 0.85-1 for Kaptive v3 and 0.09-0.94 for Kaptive v2) and for KpSC K-loci (e.g. 0.97 vs 0.82 for the non-subsampled draft assemblies for Kaptive v3 and v2, respectively), whereas the differences for the other databases were more modest (e.g. 0.99-1 vs 0.95-0.98 difference for the non-subsampled draft assemblies) (**Figure 4, Table S5**).

**Figure 4:**
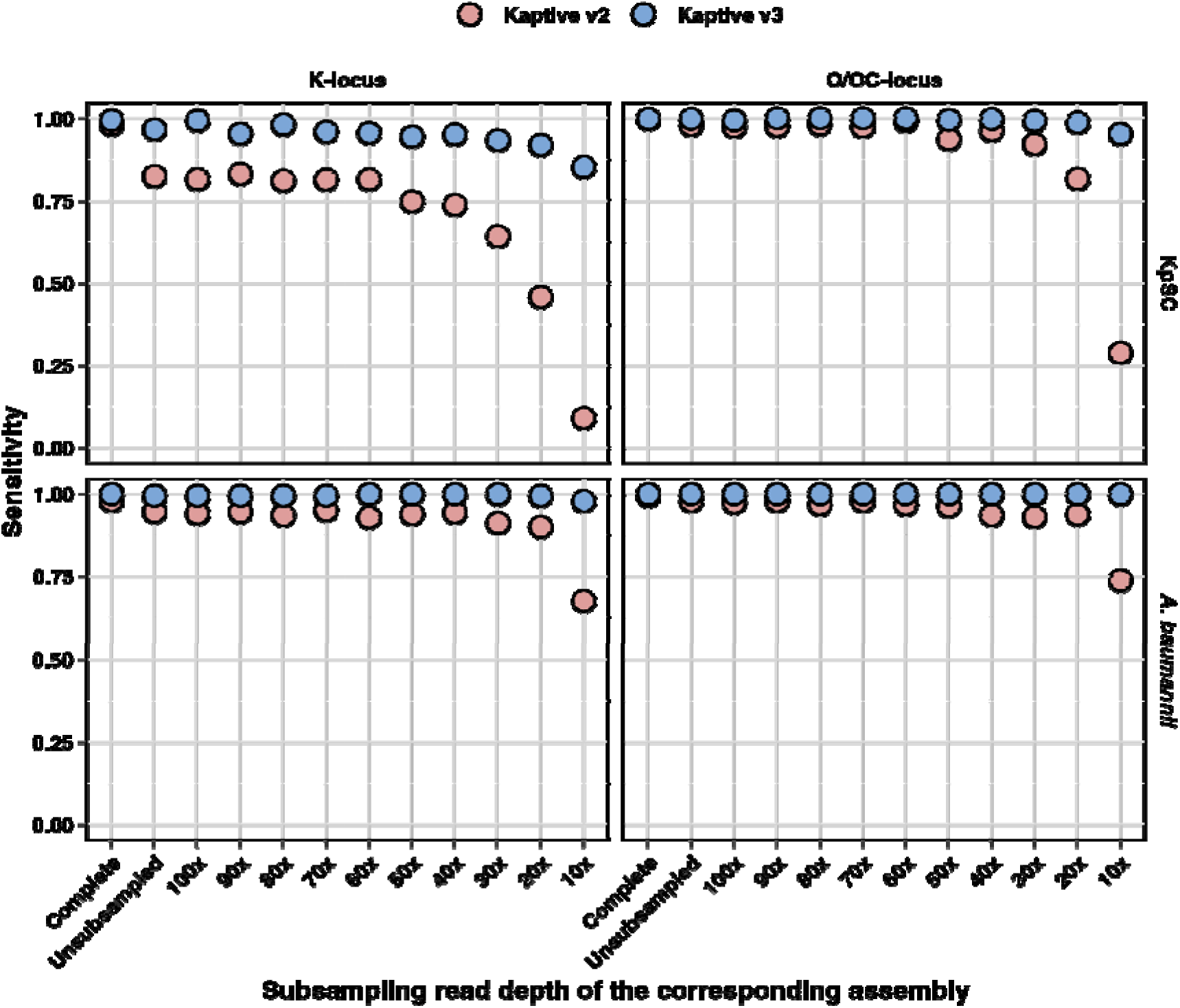
Sensitivity of Kaptive locus calls. Data are stratified by species (rows), database (columns) and assembly subsampling group, and coloured by Kaptive version as indicated. Assembly subsampling groups are arranged on the X-axis in descending order of subsampling read depth, starting with the complete (hybrid) and non-subsampled Illumina-only assemblies, then subsampling the Illumina-only reads at the stated increments.

A minority of genomes in our test dataset harboured K and O/OC-loci that were novel, which we consider genuinely untypeable. For *A. baumannii* there was a single novel OC-locus and two novel K-loci, which were correctly reported as untypeable by both versions of Kaptive. For KpSC genomes there were nine carrying novel K-loci and 10 with novel O-loci. Kaptive v2 reported between 0 and 2 of these loci typeable for each database, while Kaptive v3 reported 0-2 typeable K- and 0-3 typeable O-loci, suggesting that Kaptive v3 is slightly more likely to mistype a genuine novel locus (see **Supplementary Results** for further details).

In order to incorporate all possible outcomes into a single measure of typing performance, we calculated the percentage of assemblies reported with the optimum outcome, which we defined as: an assembly harbouring a locus with a true match in the reference database, reported correctly and marked as ‘typeable’; or an assembly harbouring a genuine novel locus, reported as ‘untypeable.’ We consider this value as a proxy for overall ‘accuracy.’ While both versions of Kaptive performed well for completed genomes (≥97% optimum outcomes for all databases), Kaptive v3 notably outperformed v2 at low assembly depths and more generally for draft assemblies typed with the KpSC K-locus database, where the percentage of optimum outcomes ranged from 7-77% for Kaptive v2 and 73-98% for Kaptive v3 (**Figure 5, Table S5**). This superior performance is primarily driven through the improvements to sensitivity, wherein Kaptive v3 is able to correctly type fragmented genome assemblies that were considered untypeable by Kaptive v2 (and also commonly misidentified). Importantly, ≥90% of all Kaptive v3 results for all assembly depths >20x (which is below standard minimum depth recommendations for Illumina genome sequencing), were considered as optimum outcomes.

**Figure 5:**
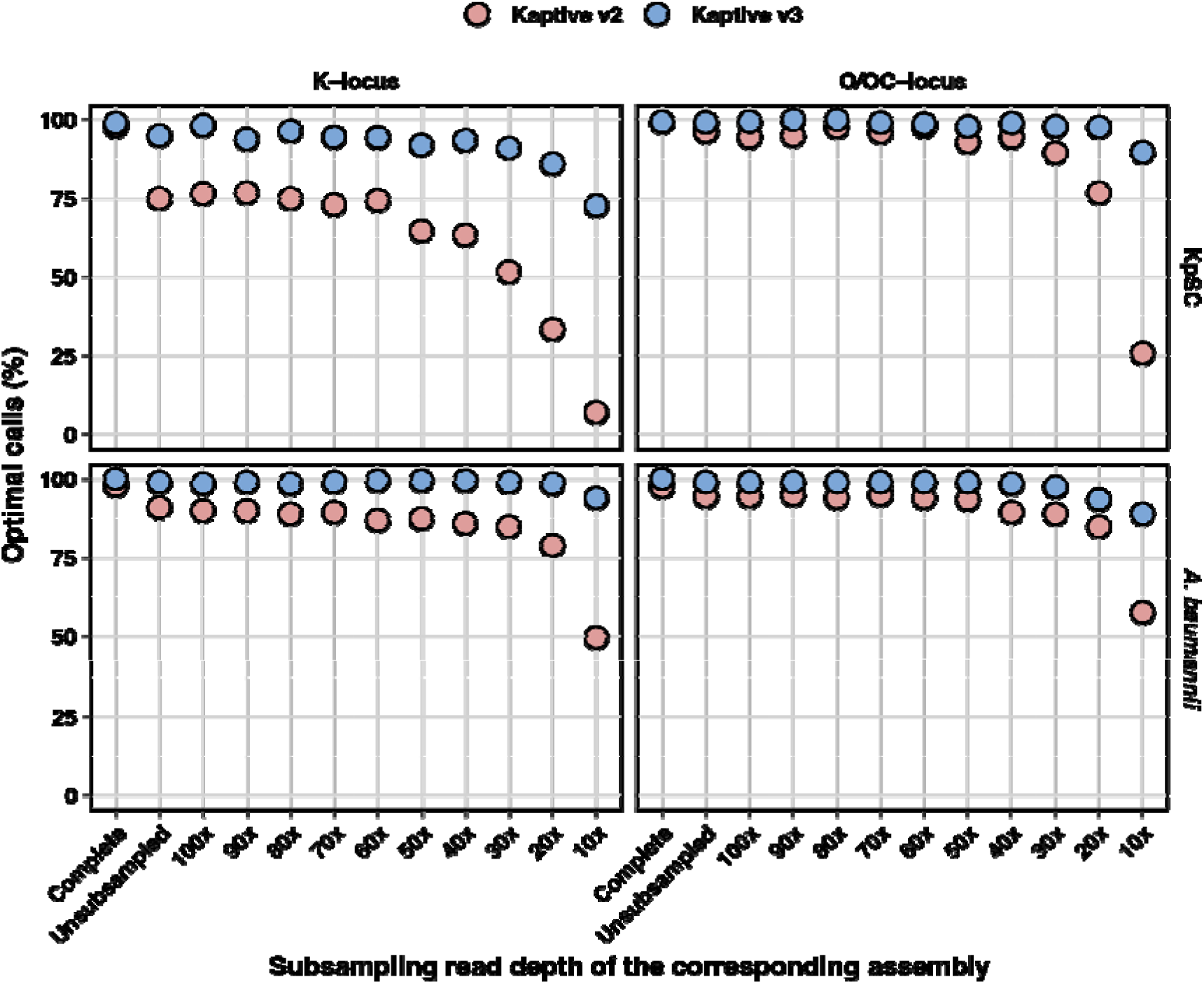
Percentage of optimal outcomes for Kaptive calls. Data are stratified by species (rows), database (columns) and assembly subsampling group, and coloured by Kaptive version as indicated. Assembly subsampling groups are arranged on the X-axis in descending order of subsampling read depth, starting with the complete (hybrid) and non-subsampled Illumina-only assemblies, then subsampling the Illumina-only reads at the stated increments.

### Kaptive v3 is Faster than Kaptive v2

Across all databases in the benchmark, Kaptive v3 outperformed Kaptive v2 in both system- and user-space, with a mean system time of 0.1 ± 0.02 vs 0.8 ± 0.5 seconds and mean user time of 1.2 ± 0.5 vs 48 ± 45 seconds (**Figure 6, Table S6**). Time in user-space reflects the external subprocesses performing the alignments and demonstrates the advantages of using minimap2 over both BLASTn and tBLASTn; whereas the decrease in system space reflects the small optimisations made to the Kaptive codebase to improve elapsed runtime across a large dataset.

**Figure 6:**
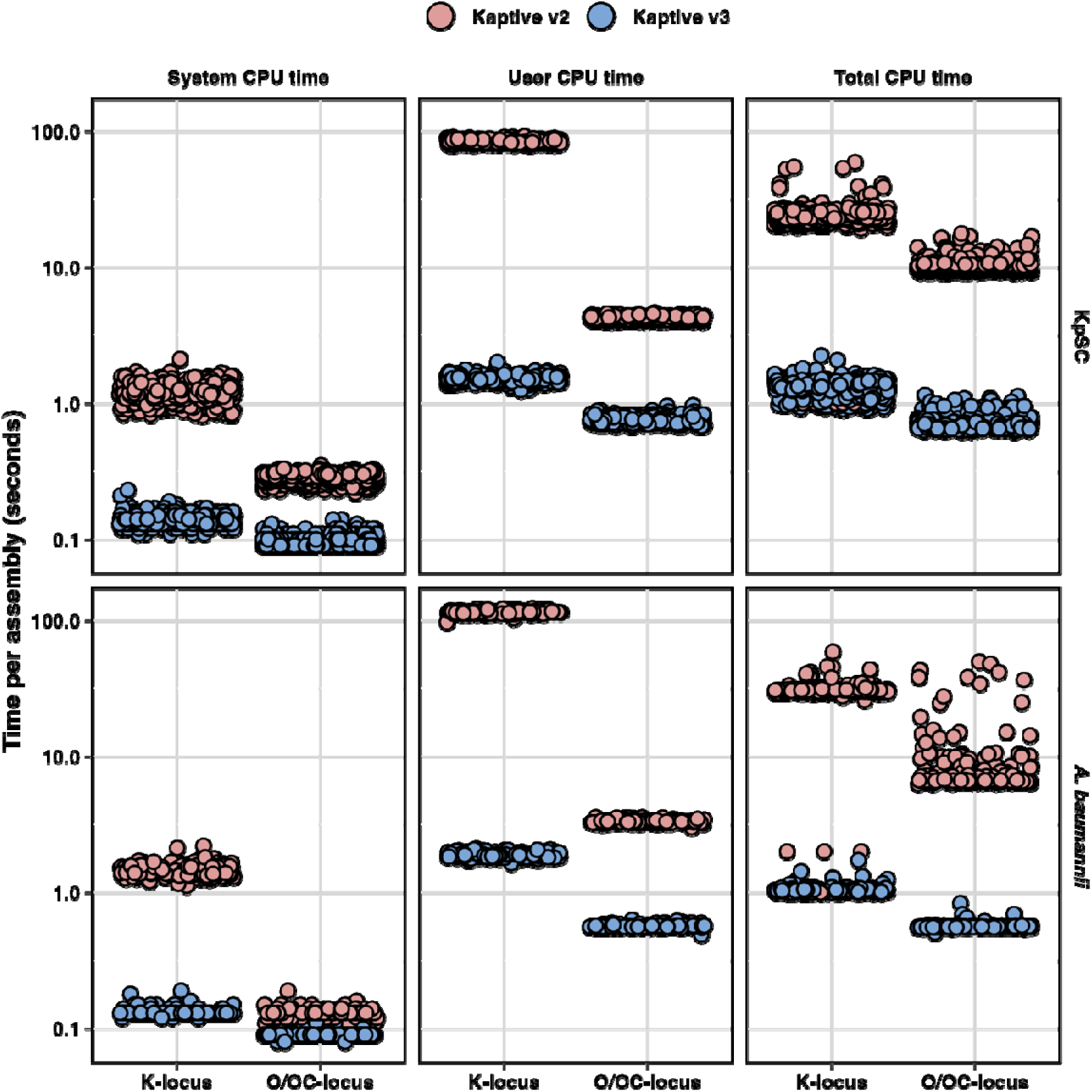
Kaptive typing speed. The three columns show system, user and total CPU time in seconds (log^10^ scaled) and the rows represent results for KpSC and A. baumannii completed assemblies stratified by database. Points represent the runtime in seconds on each assembly and are coloured by Kaptive version as indicated.

Kaptive v3 also outperformed Kaptive v2 in terms of mean total CPU time for KpSC K-locus (1.3 ± 0.2 vs 23.4 ± 5.2 seconds) and O-locus (0.7 ± 0.1 vs 10.3 ± 1.1 seconds) databases; and for *A. baumannii* K-locus (1.0 ± 0.1 vs 28.5 ± 9.0 seconds) and OC-locus (0.6 ± 0.03 vs 9.3 ± 7.3 seconds) databases (**Table S6**). The runs using the larger K-locus databases (for both species) took the longest time to execute regardless of version, especially in user-space, suggesting that execution time scales with the number of alignments performed by the respective alignment software.

Importantly when considering total elapsed CPU time, Kaptive v3 was >1 order of magnitude faster than Kaptive 2 for the KpSC K-locus (18x) and O-locus (14x) databases; and for the *A. baumannii* K-locus (16x) and OC-locus (27x) databases. This is particularly beneficial when analysing large (>1000 genomes) datasets. Whilst this performance improvement may be negated on large, distributed compute clusters (where hundreds of jobs can be run in parallel), such resources remain inaccessible to many around the world, who can now directly benefit from large-scale genomic sero-epidemiology on their desktop computers.

## Discussion

We identified key issues for *in silico* antigen typing with Kaptive and addressed them in a new version that is highly sensitive, accurate and much faster than previous versions. Notably, Kaptive v3 has higher sensitivity and results in a greater number of optimum outcomes for fragmented polysaccharide synthesis loci, facilitating accurate data extraction from many more genomes. The impact is greatest for genomes harbouring loci that are subject to sequencing drop-out due to GC divergence compared to the host chromosome (for which sequence library preparation protocols are usually optimised). These include the KpSC K-locus (a major target for sero-epidemiological analyses to inform vaccine design) [8] and are anticipated to include loci in other closely related organisms e.g. other *Klebsiella* species as well as the major pathogens *Escherichia coli* and *Enterobacter* spp. that have similar chromosomal GC content and are known to share orthologous polysaccharide synthesis genes [2]. We acknowledge that assembly fragmentation can be circumvented using long-read sequencing platforms such as Oxford Nanopore Technology’s (ONT) MinION or GridION devices, that are rapidly growing in popularity. However, we anticipate that it will be many years before these technologies become ubiquitous in research and public health laboratories, necessitating the Kaptive updates presented here. Additionally, until the recent release of ONT’s R10 sequencing chemistry, KpSC ONT-only assemblies had high rates of sequence error and untypeable Kaptive v≤2 calls due to missing genes as a result from tBLASTn mis-translation [46].

Kaptive v3’s user execution time is considerably lower than Kaptive v2, due to replacement of BLAST+ alignment subprocesses with Minimap2. The time-savings can also be attributed to a reduction in the number of alignments performed, with only the top-scoring loci fully aligned to the input assembly and the best match locus genes compared (via translation and pairwise protein alignment) in Kaptive v3. In contrast, all loci were previously aligned via BLASTn and gene content assessed with tBLASTn in Kaptive v≤2.

The ability to accurately type fragmented draft assemblies, alongside a realistic execution time for large genome collections on a personal computer, facilitates broader accessibility and greater data usability. There are currently 72,191 KpSC and 32,068 *A. baumannii* assemblies hosted on the NCBI genome database, which is increasing daily (accessed September 25th, 2024). This exponential increase in the size of publicly available datasets acts as a bottleneck for downstream analysis, and even a limiting factor for those without the compute resources to perform analysis at this scale. To further aide accessibility, including for bioinformatics-naïve users, Kaptive 3 is implemented in Kaptive-Web, and the pathogen surveillance platform Pathogenwatch, enabling thousands of users around the world to perform *in silico* antigen typing on KpSC and *A. baumannii* genome assemblies without high-performance computing resources [18,47].

We will continue to develop new Kaptive-compatible databases for other species of interest and welcome similar efforts form other teams. We performed our systematic typing performance assessments only for the Kaptive databases distributed directly with the Kaptive code; however, we expect similar performance for other databases that comprise loci distinguished by their gene content at a fixed translated sequence identity threshold, and which capture the majority of loci present in the bacterial population of interest (note ≤1.8% of genomes in our test datasets carried novel loci). Accuracy may vary for less complete databases and/or populations wherein a high number of isolates are expected to carry novel loci that may be mistyped as reference loci (**Supplementary Results**). In these cases, we recommend that users manually inspect the Kaptive output and perform confirmatory investigations for fragmented loci as well as those that are reported with large length discrepancies compared to the best match reference (i.e. which may contain novel polysaccharide processing genes that are not present in the reference database).

Finally, we would like to highlight that the accuracy estimates presented here reflect Kaptive’s capacity to detect genetic loci only. These estimates do not necessarily reflect phenotypic predictive accuracy, which is a function of locus detection *and* knowledge about the relationship between genotype and phenotype. Efforts to estimate phenotypic predictive accuracy for KpSC capsule (K) types are currently underway and will be reported elsewhere. Ongoing work will support continued improvements for both locus and phenotype prediction.

## Availability of Supporting Source Code and Requirements

Project name: Kaptive

Project home page: https://github.com/klebgenomics/Kaptive

Operating system(s): Platform independent

Programming language: Python

Other requirements: Biopython, DNA Features Viewer, minimap2

License: GNU GPL v3

## Additional Files

**Supplementary Materials** (.docx):

- Supplementary Results
- Table S1
- Figures S1-4

**Table S2 – CSV file**: **Test assemblies and Kaptive outputs.** Assembly and sequencing information (columns A-M), Kaptive version, species and database (columns N-P), Kaptive output (columns Q-AH), ground truth locus calls and Kaptive call outcomes (columns AI-AM).

**Table S3 – CSV file**: **K and O/OC-locus gene sequencing coverage and GC content.** Genome information, species and Kaptive database, Illumina read coverage (non-subsampled reads), GC content and absolute GC difference compared to the matched completed chromosome.

**Table S4 – CSV file**: **Kaptive v3 score and weighting metric combinations for the test assemblies.**

**Table S5 – CSV file**: **Performance metric summaries by assembly group and database.**

**Table S6 – CSV file**: **Kaptive compute times**

All supplementary data is available from the following figshare DOI: https://doi.org/10.6084/m9.figshare.28357046.v1

## Abbreviations

KpSC: *Klebsiella pneumoniae* Species Complex
NCBI: National Center for Biotechnology Information
ONT: Oxford Nanopore Technologies
LPS: lipopolysaccharide
CLI: Command-Line Interface

## Acknowledgements

We would like to thank Dr. Stephen Watts (University of Melbourne, https://orcid.org/0000-0001-7084-635X) for help with updating the Kaptive-Web server, and A/Prof. Johanna Kenyon (Griffith University, https://orcid.org/0000-0002-1487-6105) for help with testing Kaptive v3 with *A. baumannii* genomes.

