## Supplementary Materials for "Fast and Accurate *in silico* Antigen Typing with Kaptive 3"

### Supplementary Results

#### Kaptive typing agreement and misidentified loci

Kaptive v3 misidentified a minority of loci that were marked as ‘typeable’ results, including a single locus from a completed assembly (*A. baumannii* SAMN15646565 carrying an OCL21 IS variant, typed as OCL13), plus seven loci from non-subsampled Illumina draft assemblies (three *A. baumannii* OC loci, two K loci each for *A. baumannii* and KpSC, **Table S2**). In four of these cases the assemblies carried IS variants of reference loci that were incorrectly identified as other loci sharing a high number of gene orthologs. Kaptive v2 also misidentified the same locus from the completed assembly and for three of the seven loci from non-subsampled draft assemblies, as well as a second locus from a completed assembly and 11 additional loci from non-subsampled draft-assemblies (**Table S2**).

In contrast, many more assemblies marked as ‘untypeable’ were misidentified. Notably, the rate of agreement was as low as 13% (range 13-58% for all assembly type and database combinations with n≥20 ‘untypeable’ assemblies reported). In particular, we note that a high number of ‘untypeable’ Kaptive v2 KpSC K loci were incorrectly reported as KL107 from draft assemblies (63-81%, corresponding to up to 156 assemblies for a single assembly type and including 35 non-subsampled assemblies). This is due to assembly fragmentation resulting in BLASTn coverage bias towards the shortest locus in the KpSC K database (KL107). Unfortunately, this has led to an overrepresentation of reports of KL107 in the literature where Kaptive results were reported without consideration of confidence scores [46].

#### Kaptive typing for genuine novel loci

Among the complete and non-subsampled draft assemblies, Kaptive v3 misidentified genuine novel loci from five pairs of matched assemblies (incorrectly marked as ‘typeable’). In all cases, the ‘typeable’ confidence score criteria were met, i.e. there were no genes below the protein identity threshold, ≥50% of genes from the reported best match loci were found, and no extra genes were indicated. However, there were large length discrepancies (>2.5 – 6.6 kb) corresponding to novel genes that were not present in the reference databases and therefore were not identified as extra genes. This reflects a key caveat of the reference-based typing approach, that is expected to have minimal impact when using a reference database that contains the majority of true loci in the population of interest. However, use of databases that fail to capture the majority of loci in the underlying population will likely result in higher rates of misidentification. Future versions of Kaptive may include modules that address this issue through *de novo* annotation of contiguous loci with large length discrepancies (e.g. >1000 bp), but this will inevitably result in higher computational resource requirements and longer processing times.

### Supplementary Tables

| **Confidence** | **% Identity** | **% Coverage** | **Fragmented** | **Length difference** | **# Missing genes** | **# Extra genes** | **Reporting** |
| --- | --- | --- | --- | --- | --- | --- | --- |
| *Perfect* | 100 | 100 | No | 0 | 0 | 0 | Typeable |
| *Very high* | >=95 | >=99 | No | >=0 | 0 | 0 | Typeable |
| *High* | <95 | >=99 | No | >=0 | <=3 | 0 | Typeable |
| *Good* | <95 | >=95 ***or*** not fragmented | | >=0 | <=3 | <=1 | Typeable |
| *Low* | <95 | >=90 ***or*** not fragmented | | >=0 | <=3 | <=2 | Untypeable |
| *None* | <95 | <90 ***or*** fragmented | | >=0 | <=3 | <=3 | Untypeable |

**Table S1.** Confidence score criteria of Kaptive v≤2.

**Table S2 – CSV file**: **Test assemblies and Kaptive outputs.** Assembly and sequencing information (columns A-M), Kaptive version, species and database (columns N-P), Kaptive output (columns Q-AH), ground truth locus calls and Kaptive call outcomes (columns AI-AM).

**Table S3 – CSV file**: **K and O/OC locus gene sequencing coverage and GC content.** Genome information, species and Kaptive database, Illumina read coverage (non-subsampled reads), GC content and absolute GC difference compared to the matched completed chromosome.

**Table S4 – CSV file**: **Kaptive v3 score and weighting metric combinations for the test assemblies.**

**Table S5 – CSV file**: **Performance metric summaries by assembly group and database.**

**Table S6 – CSV file**: **Kaptive compute times**

### Supplementary Figures

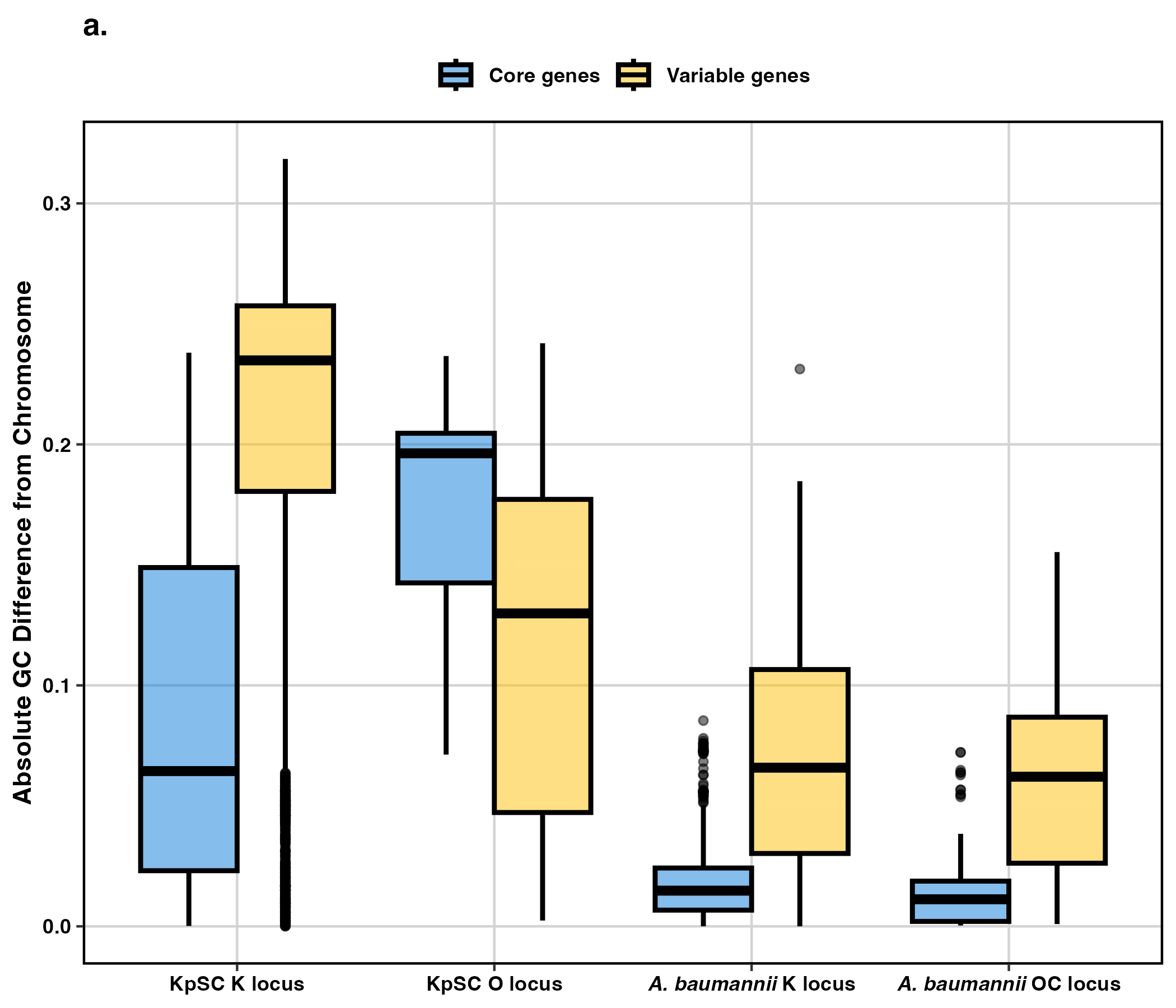

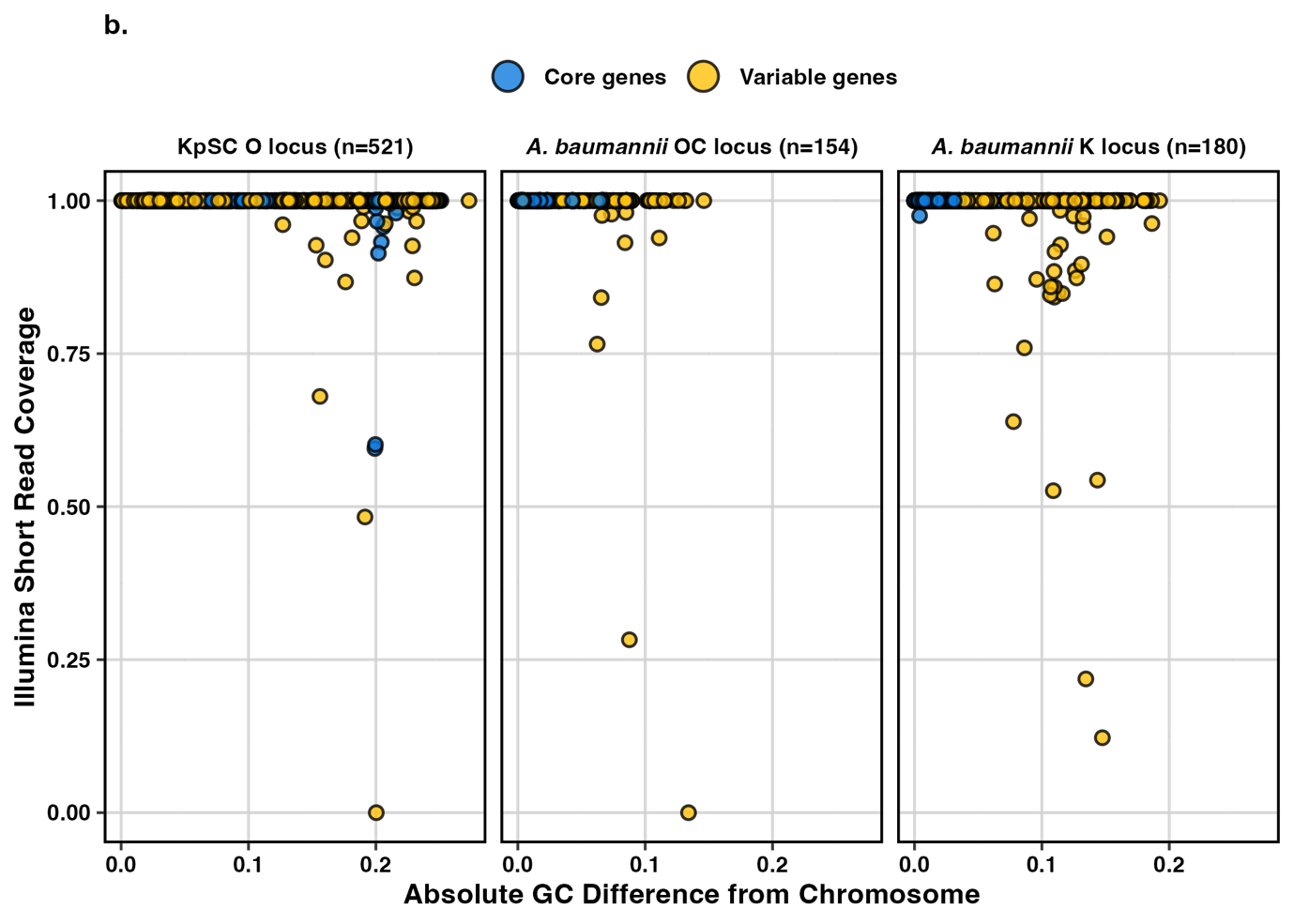

***Figure S1. Impact of GC divergence on Illumina sequencing coverage.*** *(****a****) Distributions of absolute gene GC difference from the chromosome GC value. Data are shown for core (blue) and variable (yellow) genes in the KpSC and A. baumannii K- and O/OC-locus Kaptive databases. (****b****) Illumina read coverage vs absolute GC difference of each gene compared to the chromosomal GC value. Data are shown for KpSC O-locus and A. baumannii K- and OC-locus core (blue) and variable (yellow) genes from the matched complete assemblies in the test dataset.*

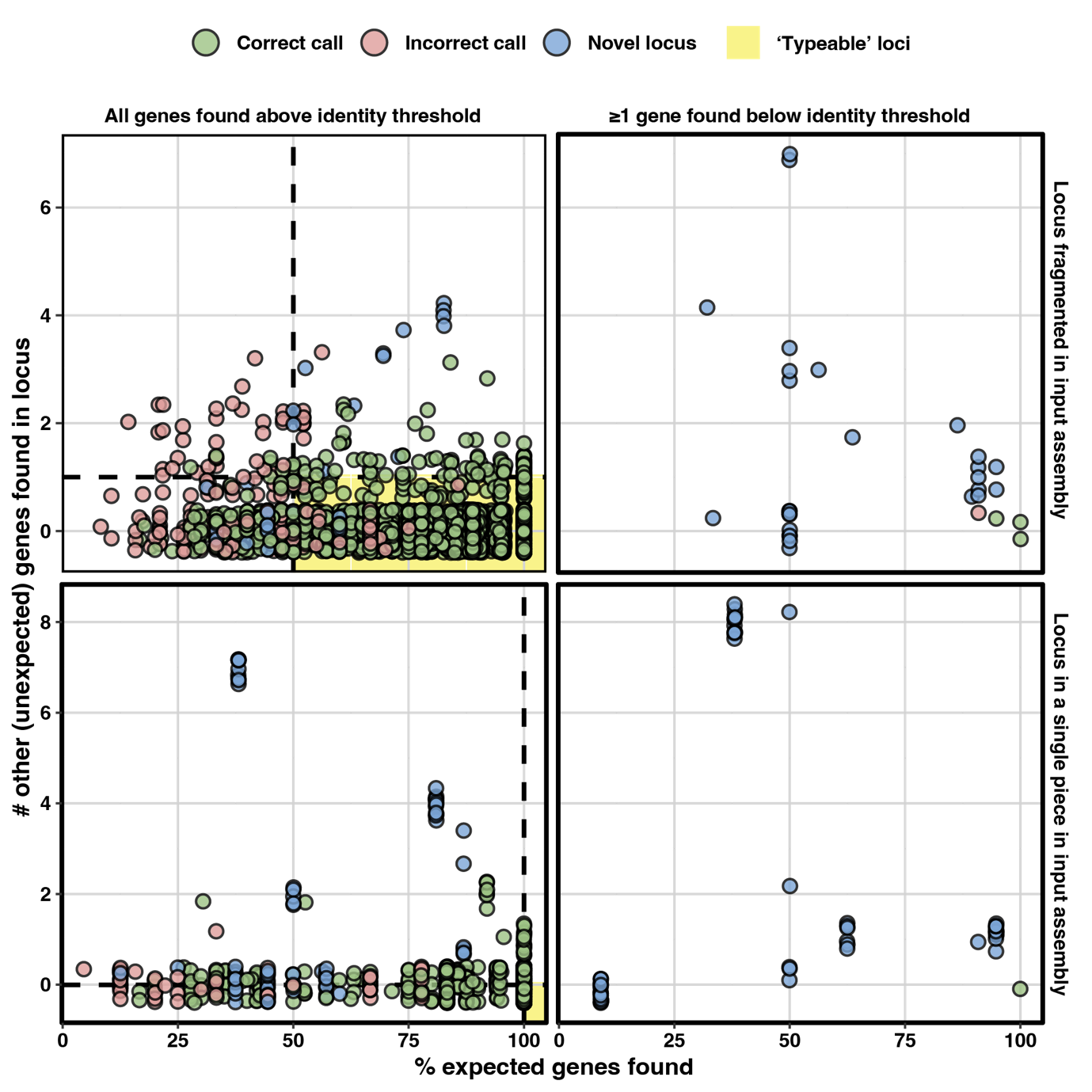

***Figure S2. Distributions of Kaptive v3 confidence scoring metrics.*** *Data represent Kaptive calls for the KpSC and A. baumannii test assemblies typed using the relevant K- and O/OC- locus databases, excluding assemblies for which the ground truth was confirmed as an IS or deletion variant (n=7,930 and n=3,392 total KpSC and A. baumannii Kaptive calls, respectively).* *The panels represent, in clockwise order from the top-left: 1) calls where the locus was fragmented in the input assembly and all genes were found above the species-specific protein identity threshold, 2) calls where the locus was fragmented in the input assembly and at least 1 gene was found below the species-specific protein identity threshold, 3) calls where the locus was found in a single contiguous piece in the input assembly and at least 1 gene was found below the species-specific protein identity threshold, 4) calls where the locus was found in a single contiguous piece in the input assembly and all genes were found above the protein identity threshold. Each dot represents an independent Kaptive locus call, coloured to indicate those that matched the ground truth (correct, green), those that did not match the ground truth (incorrect, red), and those for which the ground truth was a novel locus (blue). The Y-axis represents the number of other genes found inside the locus (i.e. those that were not expected based on the best matching reference) and the X-axis represents the percentage of expected genes found inside the locus region of the input assembly. The dotted lines represent the cutoffs implemented in Kaptive v3, and yellow shading indicates the range of values considered ‘Typeable’. Note that the individual data points are jittered to aid legibility.*

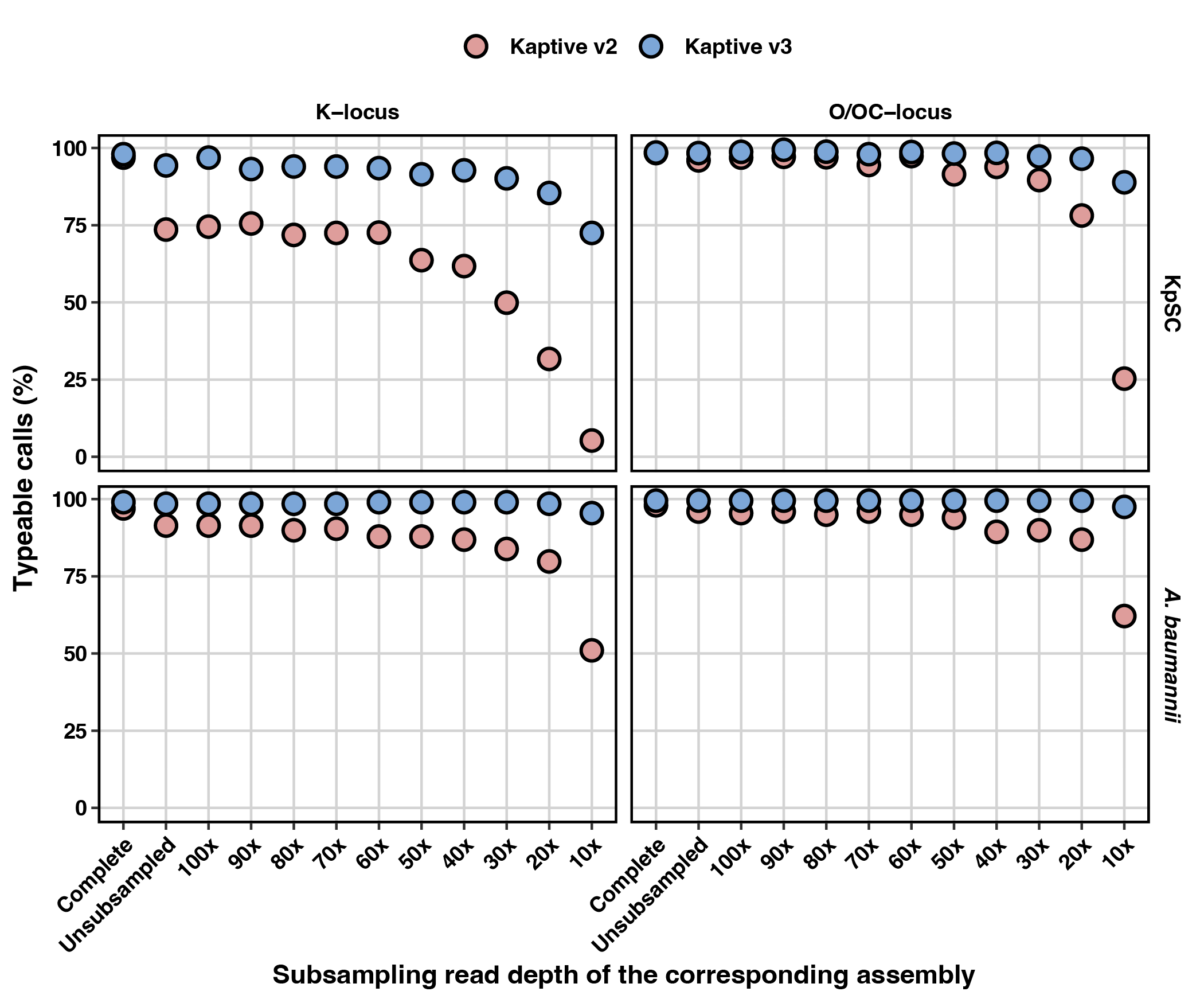

***Figure S3. Kaptive typeability.*** *Percentage of ‘typeable’ Kaptive calls, stratified by species (rows), database (columns) and assembly subsampling group, and coloured by Kaptive version as indicated. Assembly subsampling groups are arranged on the X-axis in descending order of subsampling read depth, starting with the complete (hybrid) and non-subsampled Illumina-only assemblies, then subsampling the Illumina-only reads at the stated increments.*

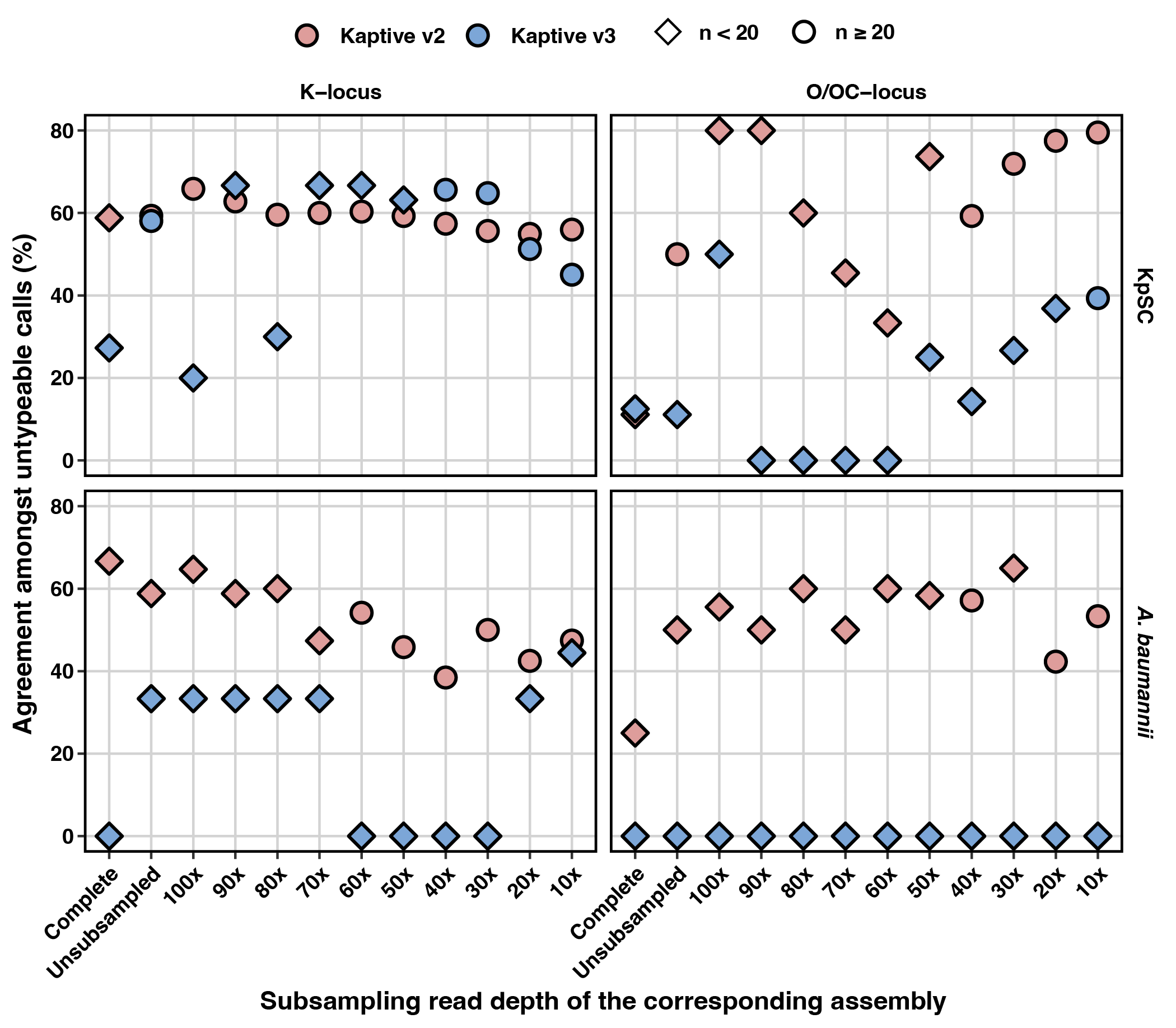

***Figure S4. Agreement among ‘untypeable’ Kaptive calls.*** *Percentage of ‘untypeable’ Kaptive locus calls that matched the ground truth. Data are stratified by species (rows), database (columns) and assembly subsampling group, and coloured by Kaptive version as indicated. Assembly subsampling groups are arranged on the X-axis in descending order of subsampling read depth, starting with the complete (hybrid) and non-subsampled Illumina-only assemblies, then subsampling the Illumina-only reads at the stated increments. Data groups for which the total number of ‘untypeable’ Kaptive calls was fewer than 20 are indicated by diamond shape.*
